# SplicingFactory – Splicing diversity analysis for transcriptome data

**DOI:** 10.1101/2021.02.03.429568

**Authors:** Benedek Dankó, Péter Szikora, Tamás Pór, Alexa Szeifert, Endre Sebestyén

## Abstract

**Motivation:** Alternative splicing contributes to the diversity of RNA found in biological samples. Current tools investigating patterns of alternative splicing check for coordinated changes in the expression or relative ratio of RNA isoforms where specific isoforms are up- or downregulated in a condition. However, the molecular process of splicing is stochastic and changes in RNA isoform diversity for a gene might arise between samples or conditions. A specific condition can be dominated by a single isoform, while multiple isoforms with similar expression levels can be present in a different condition. These changes might be the result of mutations, drug treatments or differences in the cellular or tissue environment. Here, we present a tool for the characterization and analysis of RNA isoform diversity using isoform level expression measurements.

**Results:** We developed an R package called SplicingFactory, to calculate various RNA isoform diversity metrics, and compare them across conditions. Using the package, we tested the effect of RNA-seq quantification tools, quantification uncertainty, gene expression levels, and isoform numbers on the isoform diversity calculation. We analyzed a set of CD34+ hematopoietic stem cells and myelodysplastic syndrome samples and found a set of genes whose isoform diversity change is associated with *SF3B1* mutations.

**Availability and implementation:** The SplicingFactory package is freely available under the GPL-3.0 license from Bioconductor for the Windows, MacOS and Linux operating systems (https://www.bioconductor.org/packages/release/bioc/html/SplicingFactory.html).

**Supplementary information:** Supplementary data are available at Bioinformatics online.

## Introduction

The mechanism of alternative splicing is well-known and described in most eukaryotic organisms (Lee and Rio, 2014). Alternative splicing expands the RNA repertoire of most genes, leading to changes in mRNA coding sequence or untranslated regions (UTRs). These changes might affect mRNA stability, localization or translation (Baralle and Giudice, 2017). Mis-splicing contributes to disease (Scotti and Swanson, 2015) and mutated splicing factors might act as oncoproteins or tumor-suppressors (Dvinge *et al*., 2016).

RNA-sequencing experiments regularly investigate consistent alternative splicing or mRNA isoform changes between conditions (Van den Berge *et al*., 2019). Most tools look for changes, where the ratio of mRNA isoforms or the presence of a specific alternative splicing event is coordinately increased or decreased. However, experimental evidence started to accumulate on the biological significance of gene expression variance (Eling *et al*., 2019) and more interestingly, splicing variance (Wan and Larson, 2018). Splicing variance plays a role in neurogenesis (Hattori *et al*., 2009) or innate immunity (Dong *et al*., 2012). Frequent splicing factor mutations in cancer (Seiler *et al*., 2018) might act as amplifiers of splicing variance (Wan and Larson, 2018) instead of leading to coordinated splicing changes. In turn, this increased variance might lead to large fluctuations in gene regulatory networks (GRNs), with an impact on cell fate, pathogenicity or disease penetrance. The impact of gene expression level variance on GRNs is already being investigated (Chalancon *et al*., 2012; Schuh *et al*., 2020), but similar studies do not exist for splicing regulatory networks.

A number of methods are already developed to detect changes in expression variance (Eling *et al*., 2019), but changes in splicing variance are not investigated regularly and only a few tools exist. One of the first papers in this area used Shannon-entropy to characterize splicing variance, investigating cDNA and cDNA tag libraries in 27 cancer types (Ritchie *et al*., 2008). In half of the cancers studied, they described a significant entropy gain compared to normal tissue. The RNentropy tool calculates Shannon-entropy for genes across samples to detect differential expression between any number of conditions (Zambelli *et al*., 2018). Whippet uses Shannon-entropy to detect and quantify complex alternative splicing events (Sterne-Weiler *et al*., 2018) and the authors describe that complex, high-entropy splicing events are conserved, tissue-regulated and more prevalent in various cancer types. SpliceHetero aims to characterize spliceomic intra-tumor heterogeneity (sITH) from bulk tumor RNA-sequencing (Kim *et al*., 2019). The authors used the Jensen - Shannon Divergence to characterize splice site usage differences between samples and found that increased sITH was correlated with cancer progression and worse survival. The Splice Expression Variation Analysis (SEVA) tool aims to model increased heterogeneity of splicing variants in cancer, using a rank-based multivariate statistics, comparing splice junction expression profiles between conditions (Afsari *et al*., 2018). Finally, the sQTLseekeR R package analyzes associations between genotype information and transcript relative expression. Even though the main goal of sQTLseekeR is to detect splicing quantitative trait loci (sQTLs), it can also detect splicing variance QTLs (svQTLs) (Monlong *et al*., 2014), where changes in splicing isoform diversity are associated to a genotype.

We developed the SplicingFactory R package, to facilitate the analysis of splicing isoform diversity in RNA-sequencing experiments, and investigate changes in diversity between experimental conditions with a tool that integrates into the Bioconductor package ecosystem. Our tool implements a wide range of diversity metrics, uses standardized input and output data structures and is able to process full transcriptome level expression measurements across multiple samples in a matter of minutes. Additionally, it provides individual diversity values for each gene in each sample. In contrast, the above-mentioned tools generally provide summary statistics at the sample level or between conditions. Finally, it is still largely unknow, which diversity metric might be the optimal choice for characterizing splicing variance, and we hope our tool will facilitate further research into this area.

## Methods

### Myelodysplastic syndrome dataset processing

RNA-seq datasets from two papers (Im *et al*., 2018; Pellagatti *et al*., 2018) were uniformly reprocessed. We downloaded the data from SRA (SRP133442, SRP149374) using SRA-tools (version 2.9.6). For sample ids and sample status see Supplementary Table 1. We checked sample quality using FastQC (version 0.11.9) and MultiQC (version 1.9). In the case of the SRP149374 data, we merged the downloaded fastq files into a single file, dropping an outlier run from each sample with different read length from the rest. We aligned RNA-seq reads to the GRCh38 reference genome without alternative contigs, using STAR (version 2.7.2b) in 2-pass mapping mode. We quantified transcript level expression using Kallisto (version 0.46.2) and Salmon (version 1.2.1) and the full GENCODE v34 transcriptome annotation. Kallisto was run with 100 bootstraps, the --bias parameter, and the output format was set to plain text. We generated the Kallisto index based on the full GENCODE v34 transcript fasta file. Salmon was run both in alignment-based using the STAR alignments and mapping-based mode (referred to as Salmon-SAF in the text) using a decoy-aware transcriptome index. We generated the Salmon transcriptome index using the GENCODE v34 transcript fasta file, combined with the full GRCh38 reference genome without alternative contigs as the decoy sequence. In both cases we run the tool with 100 bootstraps, the unstranded paired-end library option and the additional --seqBias and --gcBias parameters. The mutation status of samples was defined based on the original papers’ supplementary material. We calculated the various diversity metrics across samples and genes using the Kallisto, Salmon or Salmon-SAF Transcript Per Million (TPM) expression estimates.

### Comparison of Kallisto, Salmon and Salmon-SAF results

In order to compare the consistency of the diversity metrics calculated from the three different sources (Kallisto, Salmon, Salmon-SAF) we selected the 17 control samples from the SRP133442 dataset and calculated the normalized naive entropy and Gini-index for all genes. For the three pairwise comparison (Kallisto – Salmon, Kallisto – Salmon-SAF, Salmon – Salmon-SAF) and each sample we selected all genes with non-NA diversity values and calculated their Spearman correlation.

### Analyzing uncertainty of estimated isoform abundances

To analyze the effect of expression estimation uncertainty on diversity metrics, we processed the Kallisto, Salmon and Salmon-SAF bootstrap data of the 17 control samples from the SRP133442 dataset. In the case of Salmon and Salmon-SAF, we first converted the binary bootstrap data to text format using the script ConvertBootstrapsToTSV.py from the GitHub page of Salmon (https://github.com/COMBINE-lab/salmon), and converted the estimated read counts to TPM values. Using the bootstrap data, we calculated all available diversity metrics for all genes from all samples and bootstraps. Finally, we selected the (Hay *et al*., 2018) CD34+ hematopoietic stem marker genes and checked the consistency of the bootstrapped expression values and the calculated diversity metrics of these genes in one specific good quality control sample (SRR6781226).

### Filtering low abundance transcripts

To analyze the effect of filtering low abundance transcripts on the calculated diversity metrics, we used the 17 control samples from the SRP133442 dataset. First, we filtered the Kallisto, Salmon and Salmon-SAF transcript-level expression data using the following criteria. For each filtering step, we kept only those transcripts that had a relative TPM abundance level compared to their gene’s total expression over a pre-defined threshold across all 17 samples. The applied relative abundance thresholds were 1, 2, 5, 7, 10, 15 and 20 percentages. This way, we created 3 × 7 abundance tables with different transcript compositions. We then calculated all 7 diversity metrics available in the package. Finally, we calculated the mean Spearman correlation of the diversity values of the filtered and non-filtered data of the 17 samples and calculated 95% confidence intervals, using R (version 4.0.2) and the bootstrap package (version 1.3-25) using 500 bootstrap replicates.

### Performance benchmarks

The performance benchmark was done using a single core on a server with Intel^®^ Xeon^®^ Gold 4118 2.30 GHz CPU type, a total of 157 GB memory and Ubuntu 18.04.5 LTS operating system. We used the Salmon-based quantification for the benchmarks and tested both the diversity and difference calculation steps using the samples from the two MDS datasets. We calculated elapsed time in seconds and maximum memory used in megabytes for the diversity calculation step as follows. Using a fixed number of 60669 starting genes, we increased the sample number from 10 to 130 using steps of 10, 20, 40, 60, 80, 100 and 130 for the same calculations. To test the difference calculation step, we used the same increasing sample numbers as in the diversity calculation performance benchmarks and tracked resource usage with the same bash shell command.

### Comparison to other tools

We used SpliceHetero (version 1.0), SEVA (available in the GSReg R library version 1.24.0) and Whippet (version 1.6.1) for comparison, analyzing the Salmon-based quantification results of the 17 control and 17 MDS samples from the SRP133442 dataset.

We ran SpliceHetero with parameters -slb True and -prn 17. To prepare the input data, we first filtered the STAR splice junction files, keeping only those splice junctions that appear in the STAR index sjdbList.fromGTF.out.tab file and had at least one uniquely mapping read.

Then, we converted the filtered STAR junction files to BED6 format, and sorted the output file. Finally, we ran SpliceHetero on these sorted BED files and we annotated the output splice junction level entropy values with the GENCODE v34 transcriptome annotation. Finally, we averaged these entropy values across genes and used these values for further comparison.

We ran SEVA in RStudio Server (R version 4.0.2) using the GSReg SEVA function from the GSReg library with the parameter single.strand.genes.only set to FALSE and used the Bioconductor annotation database TxDb.Hsapiens.UCSC.hg38.knownGene (version 3.10.0) for gene annotation. To prepare the input data, we first performed RPM (Reads Per Million) normalization on the STAR splice junction files considering uniquely mapping reads only. We used the SEVA gene-level E1 and E2 variance estimates for further comparison.

We ran Whippet with default parameters. To prepare the input data, we converted the raw fastq files to have only a single “+” character on every third line. Then, using Whippet’s whippet-index.jl script, we created an index file based on the GENCODE v34 transcriptome annotation gtf file, and ran Whippet’s quantification algorithm with the whippet-quant.jl script on the modified fastq files with the additional --biascorrect parameter. Finally, we compared the MDS and control samples using Whippet’s whippet-delta.jl script. Finally, we averaged Whippet’s splicing event level entropy estimates across genes and used these values for further comparison.

### SF3B1 differential diversity and enrichment analysis

To demonstrate diversity changes between sample groups, we analyzed the SRP149374 dataset. We compared the MDS samples without any known mutation to MDS samples with known *SF3B1* somatic mutations based on the original paper describing the dataset (Pellagatti *et al*., 2018). We calculated the normalized naive-entropy, normalized Laplace-entropy, and Gini-index using the Salmon-based quantification, then compared the *SF3B1* mutated and *SF3B1* wild-type samples using Wilcoxon-test and performed P-value adjustment with the Benjamini-Hochberg method.

We also performed enrichment analysis using the significant genes (|mean difference| > 0.1 and adjusted P-value < 0.05) separately with either a mean diversity increase or decrease between the *SF3B1* mutant and *SF3B1* wild-type groups. We used the bone marrow marker gene sets originating from (Hay *et al*., 2018) and downloaded from MSigDB (Liberzon *et al*., 2015). We performed a Fisher’s exact test using R (version 4.0.2) with the *alternative=”greater”* option. Moreover, we performed P-value adjustment with the Benjamini-Hochberg method.

## Implementation

The splicing isoform diversity analysis works as a two-step process: 1) the package calculates a diversity value for each gene in each sample, using splicing isoform expression values, and 2) calculates differential diversity results between conditions. Diversity values from 1) can be used independently from step 2) for custom downstream analyses or visualizations.

### Input data structure

The package can process R matrices and data frames with expression values, assay data from the SummarizedExperiment Bioconductor object, data from the DGEList object of edgeR (Robinson *et al*., 2010), or the output of tximport (Soneson *et al*., 2016). The package requires that samples are specified as columns and transcript level expression values are specified as rows. Additionally, it needs a vector of genes used to aggregate and analyze the splicing isoform level data, and a vector of sample categories used to calculate differential diversity. All of the data structures might contain RNA-sequencing read counts, RPKM, FPKM or TPM values.

While the package can process any kind of numeric value, used to measure expression levels, we recommend TPM or a similar length normalized value. Read count values not normalized for transcript isoform length, and diversity values based on them might be misleading. For example, a gene with three transcript isoforms, and lengths of 100, 100, and 1000, and read counts of 20, 20 and 200 for each of them is detected in an experiment. Simply using the read counts to calculate proportions will lead to the values of 0.083, 0.083 and 0.83, and to the conclusion that we have a single dominant isoform based on the diversity value. However, the third isoform is 10 times longer than the other two, leading to a larger number of reads originating from it just by chance. Normalizing for isoform length will lead to the same 0.33 proportion for all isoforms, therefore no dominant isoform and a very different diversity value.

### Diversity calculation

As a first step, the package calculates diversity values using the calculate_diversity() function for each gene and each sample. Multiple diversity measures are implemented, including the Shannon-entropy, Laplace-entropy, Gini-index, Simpson-index and the inverse Simpson-index.

Shannon-entropy is a classic measure of uncertainty in information theory, in our case, ranging from 0 to log_2_(isoform-number) for a gene. As the maximum value of Shannon-entropy depends on the number of splicing isoforms for a gene, we implemented a normalized Shannon-entropy, that ranges between 0 and 1. This makes it possible to compare entropy values of genes with different number of isoforms. A 0 Shannon-entropy means a single splicing isoform is expressed from a gene, while 1 means all isoforms are evenly expressed, with no dominant isoform. The package can also calculate Laplace-entropy, a Bayesian estimate of the Shannon-entropy, where a pseudo-count of 1 is added to the isoform categories for each sample.

The Gini-index is originally intended to represent income inequality in economics. It ranges between 0 and 1, where 0 means complete equality, *i*.*e*. all isoforms have the same expression, while 1 means complete inequality, with only a single isoform being expressed. The Simpson-index is a measure of diversity originally used in ecology to quantify species diversity. A 0 Simpson-index means low diversity, *i*.*e*. one dominant isoform, while 1 means high diversity, where all isoforms have the same expression. Similar to the non-normalized Shannon- and Laplace-entropy, the maximal value of the Simpson-index depends on the isoform number. The inverse Simpson-index starts at value 1, and higher values mean greater isoform diversity, the maximal value also depending on isoform number.

The calculate_diversity() function returns a SummarizedExperiment object, that contains the gene level splicing diversity values, together with gene names, sample ids, and metadata information, including the method used, and if normalization was applied. The function removes genes with a single isoform and adds an NA value for genes where the expression of all isoforms is 0 in a specific sample, and a meaningful diversity value is impossible to calculate.

### Differential diversity calculation

Users can calculate splicing diversity changes between two conditions using the calculate_difference() function. Accepted input formats are R data frames or a SummarizedExperiment object. In the case of a data frame, gene names must be present in the first column, and splicing diversity values in all additional columns. Differences and log_2_ fold changes in diversity can be calculated using the mean or median values across conditions. The function returns the mean or median for both conditions, the difference of means or medians, and their log_2_ fold change.

Caution must be taken when choosing and interpreting a difference metric for a specific diversity type. The normalized Shannon-entropy, the normalized Laplace-entropy and the Simpson-index are all bounded in [0, 1] and as transcriptomic diversity increases their value also increases. In contrast, while the Gini-index is also bounded in [0, 1], its value decreases as the transcriptomic diversity increases. Finally, the inverse Simpson-index and the non-normalized Shannon and Laplace entropy are not bounded in [0, 1]. The inverse Simpson index starts at 1, while the non-normalized entropies start at 0, and their maximal value depends on the isoform number. Therefore, we recommend using the mean or median difference for values bounded in [0, 1] not to over-emphasize relatively small changes magnified by the log_2_ scale ratio. However, in the case of values not bounded in [0, 1] we suggest using the log_2_ fold-change.

Statistical significance of the changes can be assessed using a Wilcoxon-test or sample label shuffling and the function returns the p-values, together with the FDR corrected ones. It automatically excludes genes from the significance analysis, where some of the sample diversity values are missing and the total number of samples is insufficient for significance calculation.

## Results

### Association of diversity metrics with gene expression and isoform number

Using the reprocessed RNA-seq datasets, we investigated the effect of gene expression and isoform number on the various diversity metrics. **Figure 1** shows all 7 implemented diversity metrics. As expected, the Gini-index shows an opposing pattern compared to all others. The non-normalized naive- and Laplace-entropy, besides the Simpson-index and the inverse Simpson-index shows a clear association with isoform number, as the maximum value of the metric is determined by the number of isoforms. This association is not seen for the other three metrics. The horizontal patterns present on the non-normalized Laplace-entropy panel are the result of the +1 pseudocount of Laplace-entropy, that exaggerates the maximum values for different isoform numbers. Finally, the strongly biased pattern for the normalized Laplace-entropy is again the result of the +1 pseudocount, as all genes with only zero or very low isoform expression levels are given an expression of ∼1, leading to the maximum possible entropy of 1.

**Figure 1:**
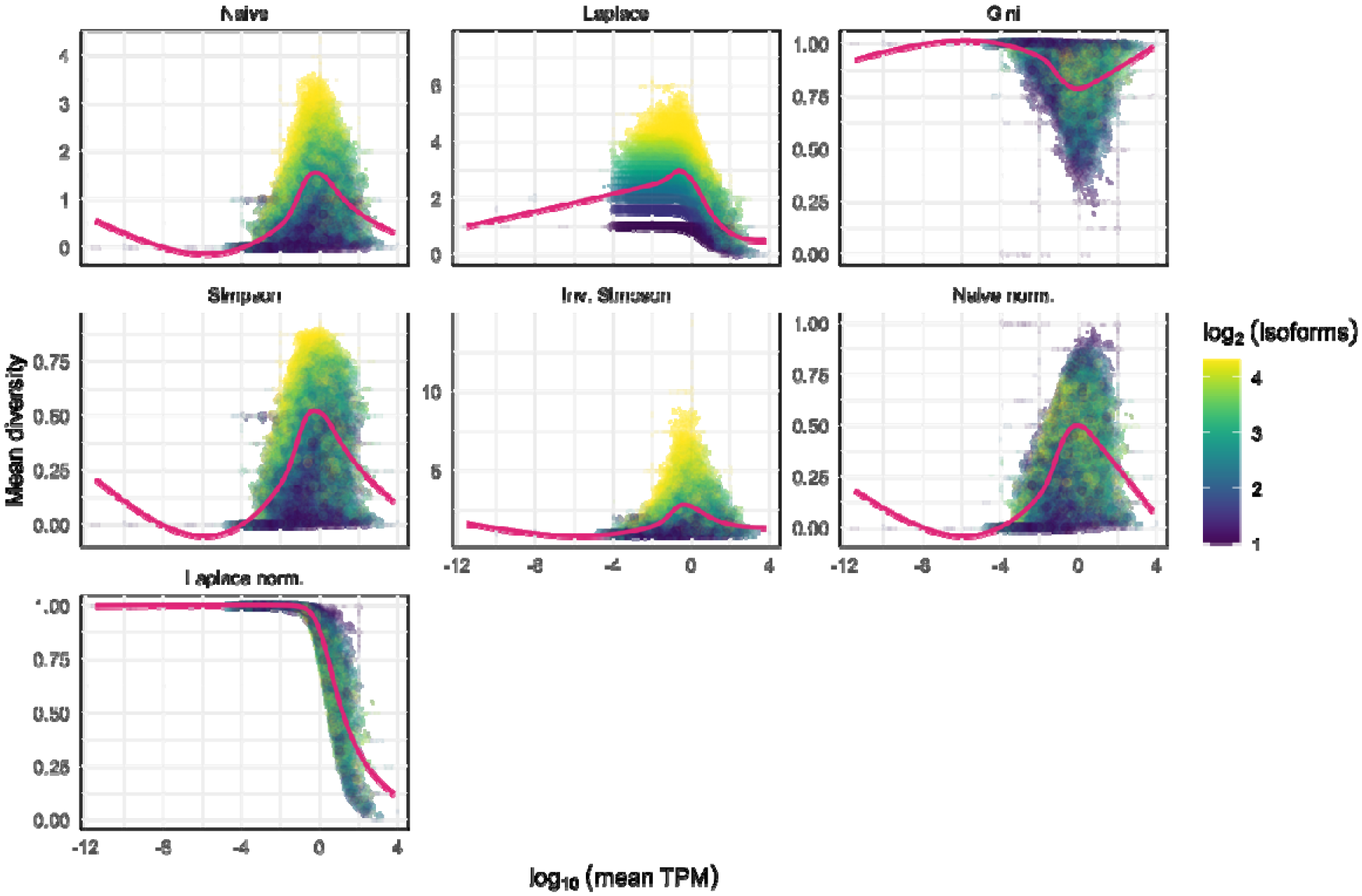
Correlation of various diversity metrics with gene expression and isoform number. Each dot represents a gene with at least two isoforms and at least one protein coding isoform. The x axis shows the log_10_(mean TPM) expression across the 17 control samples from the SRP133442 dataset, while the y axis shows a given mean diversity metric across the same samples. The color of the dots shows the log_2_(isoform number) for the gene. Gene with more than 20 isoforms were assigned the value 20. The purple line shows the smoothed conditional mean of the data using a generalized additive model. Naive – non-normalized naive entropy, Laplace – non-normalized Laplace-entropy, Gini – Gini-index, Simpson – Simpson-index, Inv. Simpson – inverse Simpson-index, Naive norm – normalized naive entropy, Laplace norm: normalized Laplace-entropy.

### Influence of different quantification tools on diversity metrics

Using the same datasets, we assessed the effect of different transcript quantification tools on diversity metrics. We calculated the naive normalized entropy and the Gini-index for all genes across all samples and calculated the Spearman-correlation of these values between methods for all samples. **Figure 2A** shows the distribution of the correlations. The Salmon-SAF – Kallisto comparison shows the highest correlation, while Salmon in alignment-based mode, using the STAR alignment output leads to significantly different diversity metrics as can be seen on the Salmon – Kallisto comparisons. As Salmon in alignment-based mode uses information from reads aligned to the full genome, it is not surprising that results are different from the other two methods, not using read alignment information. Based on these results, transcript level expression quantification tools might have a significant effect on diversity calculations.

**Figure 2:**
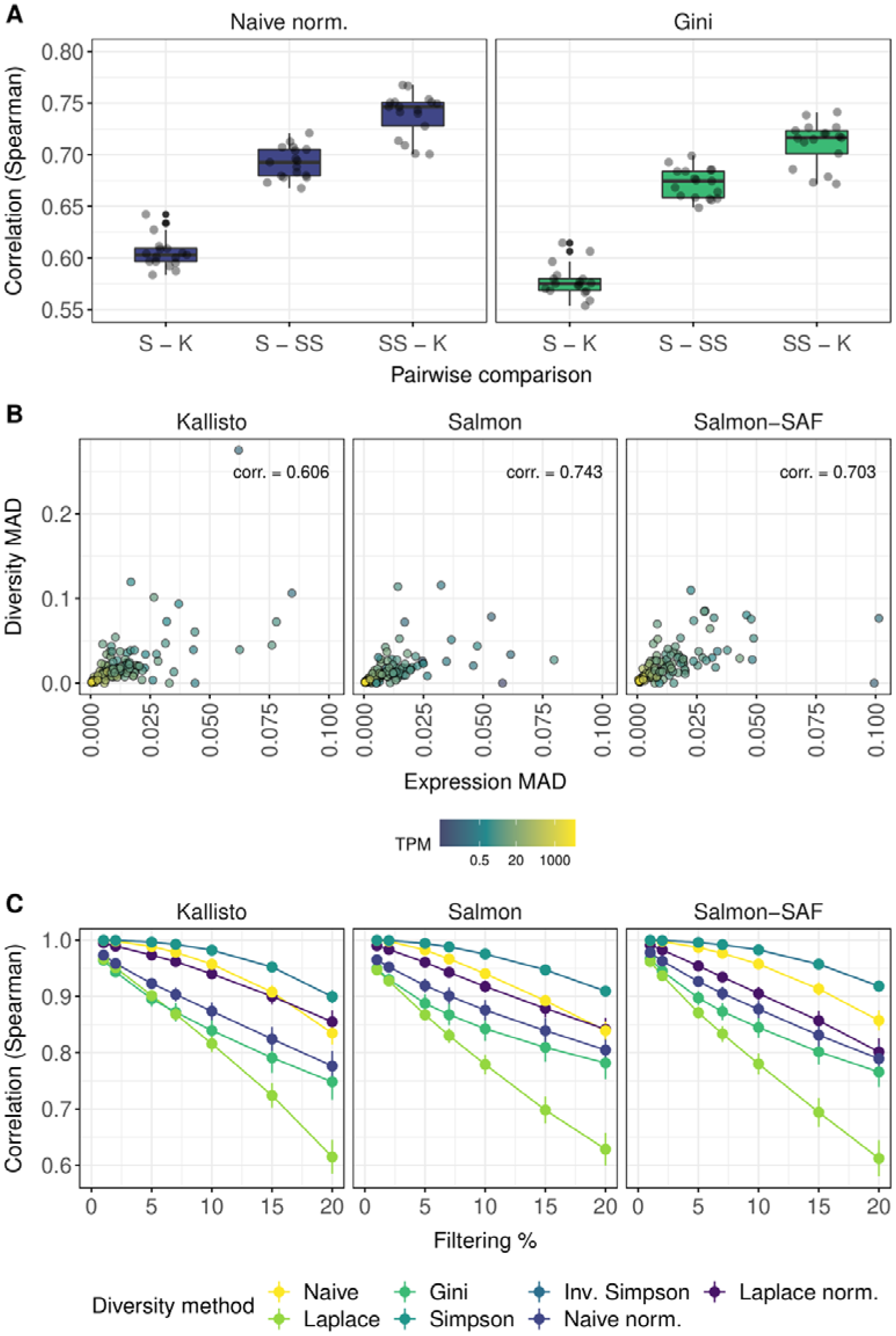
Various factors influencing diversity metrics. **A**) Consistency of diversity metrics using the Kallisto, Salmon and Salmon-SAF transcript quantification methods. Each boxplot shows the Spearman correlation values for a specific pairwise comparison in the 17 control samples from the SRP133442 dataset, showing two different diversity metrics. K – Kallisto, S – Salmon, SS – Salmon SAF. **B**) Influence of expression estimation uncertainty on diversity calculation. The x axis shows the expression median absolute deviation (MAD), while the y axis shows the naive normalized entropy MAD of selected marker genes in a single sample. Dot color shows the original non-bootstrap expression estimates for the gene. The corr. value on the panels is the Spearman-correlation of the expression MAD and the diversity MAD. **C**) Influence of filtering low-expression transcripts before diversity calculation. The x axis shows the % criteria used for filtering transcripts (see Methods), while the y axis shows the Spearman correlation of the diversity metrics using filtered results and the original unfiltered results. Vertical lines at the dots shows bootstrap replicate confidence interval.

### Effect of estimated isoform abundance uncertainty

To further investigate how quantification tools, and transcript expression levels influence diversity calculation, we selected a set of CD34+ hematopoietic stem cell marker genes from the (Hay *et al*., 2018) dataset and a single sample for analysis (see Methods). After calculating 100 bootstrap expression estimates for all three tools, we calculated the naive normalized entropy for each bootstrap, and calculated their median absolute deviation (MAD). Additionally, we calculated the MAD for the gene-level TPM expression values. We show the results on **Figure 2B**. Additionally, we also calculated the Spearman-correlation between the diversity MAD and the expression MAD for all three tools across all genes. As can be seen, a higher expression MAD leads to higher diversity MAD values. As diversity values are based on the transcript level expression estimates, uncertainty in transcript expression estimation leads to uncertainty in diversity calculation. However, genes with a higher overall expression level, based on the non-bootstrap value, tend to have lower MAD both for expression and diversity.

### Effect of filtering low-expression isoforms

Finally, we investigated the effect of removing an increasing number of transcripts with low expression from our analysis. **Figure 2C** shows the results, where the non-normalized Laplace entropy was affected the most, regardless of the quantification method used. In other cases, removing low expression transcripts from the analysis changed the diversity values somewhat, and the correlation with the original dataset decreased, but they were still in the range of 0.75 – 0.99 in all cases. Therefore, it might be useful to remove transcripts with very low or zero expression at the start of the analysis, and this would not compromise results significantly. This is especially important for genes with a large number of transcripts, and low expression, where the number of reads from the sequencing experiment is insufficient to give a confident expression and therefore confident diversity value.

### Performance benchmarks

Using the Salmon expression estimates from the reprocessed datasets, we carried out a benchmark, investigating the total runtime and memory consumption of SplicingFactory using a single CPU core. **Figure 3A** shows the maximum memory usage of the package, when calculating a specific diversity metric or also calculating differential diversity between conditions. As we increased the sample number, memory usage also increased linearly, with a maximum of 2500 megabytes for 130 samples. However, even the maximum memory usage is well within the range of a regular desktop computer and the package is not limited by memory limitations.

**Figure 3:**
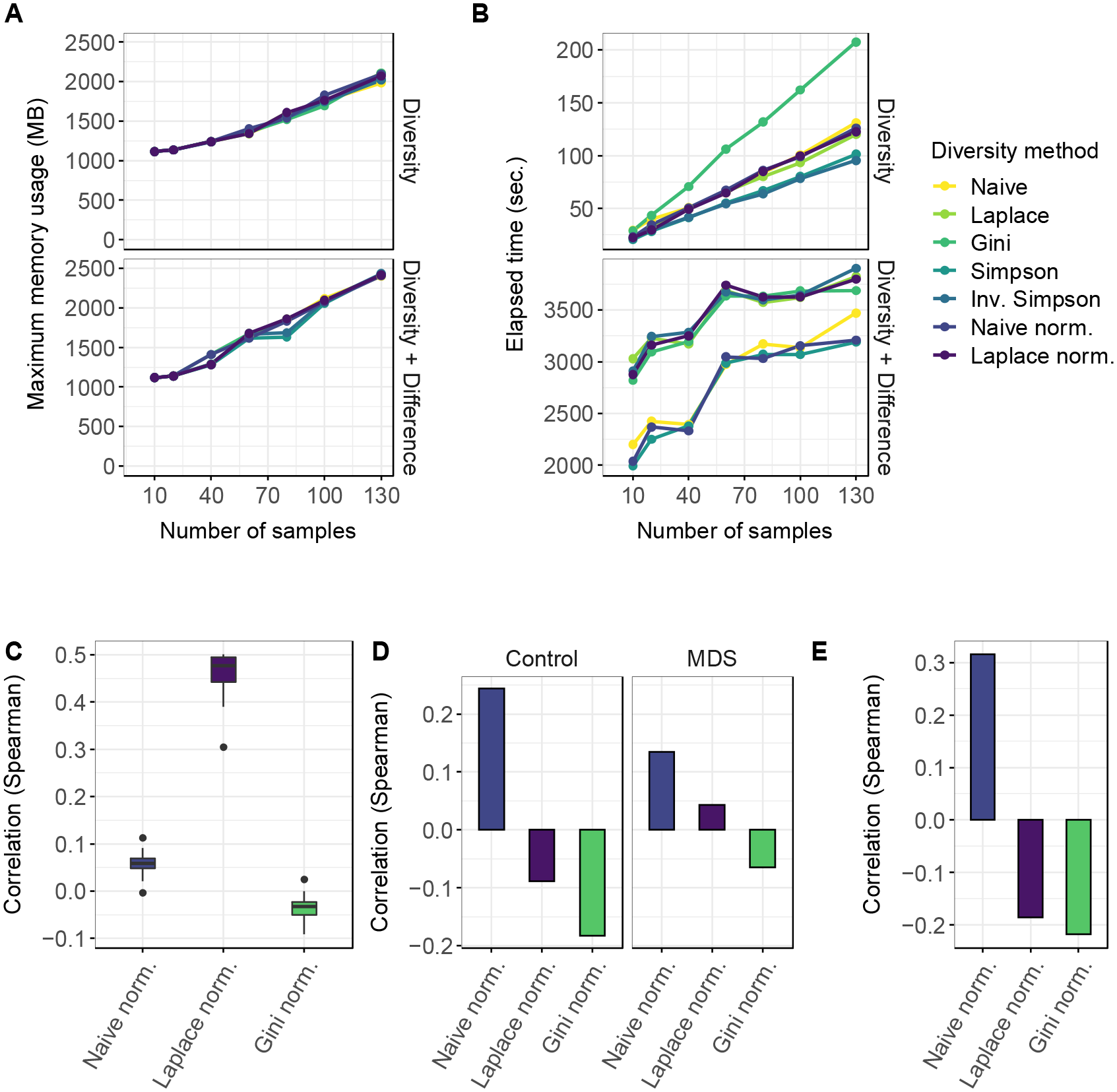
Performance benchmarks and comparison of SplicingFactory to other tools. **A**) Memory usage of SplicingFactory with increasing sample number for all diversity metrics while calculating the diversity values or also calculating differential diversity between sample groups. The x axis shows the increasing number of samples used, while the y axis shows the maximum amount of memory used. **B**) Total elapsed time for calculating the diversity values or also calculating differential diversity between sample groups for all diversity metrics. The x axis shows the increasing number of samples used, while the y axis shows the total elapsed time for the calculation. **C**) Spearman correlation of SpliceHetero entropy values with 3 different diversity metrics and Salmon expression estimates for the 17 MDS samples from the SRP133442 dataset. **D**) Spearman-correlation of average modified variance values (E1 and E2) as calculated by the GSReg.SEVA function and average diversity values for 3 different diversity metrics for the 17 control and MDS samples from the SRP133442 dataset. **E**) Spearman correlation of Whippet entropy and average diversity values for 3 different diversity metrics for the 17 control and MDS samples from the SRP133442 dataset.

We also tested the total running time for the same setup, and results are shown on **Figure 3B**. Calculating the Gini-index takes the most time, when considering only the diversity calculation step, but even this calculation takes only 3 minutes. The additional differential diversity calculation peaks at almost an hour for 130 samples using some of the diversity metrics. Even with an increased running time during differential diversity calculation, our tool is able to process the expression estimates of a full transcriptome annotation across more than 100 samples. This requires only a limited amount of resources and approximately an hour using a single core, making it feasible to run the analysis on any up-to-date desktop PC or laptop.

### Comparison to other tools

In addition to the performance benchmark, we compared the results of our tool to three additional tools characterizing splicing diversity. Using the Salmon expression estimates for the 17 control or 17 MDS samples from the SRP133442 dataset, we focused on comparing the actual gene and sample level diversity metrics of SpliceHetero, SEVA and Whippet, if available to our tool.

We used the gene-level averaged entropy estimates of SpliceHetero of the MDS samples and calculated their correlation with our diversity metrics for the same samples. Results are shown on **Figure 3C**. Both the normalized naive entropy and the Gini-index show very low correlation, between −0.1 – 0.1. Correlation of the normalized Laplace-entropy is much higher, between 0.3 and 0.5. Unfortunately, SpliceHetero does not provide entropy values for samples defined as a control set. Control samples are only used as a baseline, against with another set of samples are compared, to see if entropy changed. Additionally, there is no significance testing of results, only the change in entropy values are provided. In the case of SEVA, we compared the E1 and E2 variance metrics calculated for the control and MDS samples, respectively. As shown on **Figure 3D**, correlations for all three diversity metrics range between −0.2 – 0.2, with control sample values being slightly higher. Based on these results, values calculated by SEVA and our package seem to be significantly different, without much overlap or correlation at the gene level. Finally, we compared Whippet’s splicing event level entropy values averaged across genes to our diversity metrics. Similar to the previous comparisons, correlations range between −0.2 – 0.3 for all three diversity metrics, as shown on **Figure 3E**.

### Association of splicing factor mutations with changing diversity

Finally, using the Salmon expression estimate based results, we investigated differences in diversity between splicing factor mutated and non-mutated MDS samples. We compared the *SF3B1* mutated and wild-type samples from the SRP149374 dataset and results are shown on **Figure 4A**. Using the normalized naive-entropy, normalized Laplace-entropy and Gini-index metrics, we found 97, 154 and 27 genes with significant changes, respectively. Based on the normalized naive-entropy we detected 94 genes where diversity increased, and 60 where diversity decreased. In the case of normalized Laplace-entropy and Gini-index, we detected 28 + 69 and 15 + 12 genes, respectively. Overall, we found 242 unique genes across the three comparisons. Six genes were detected by all three metrics, 24 by two of them, and 212 by a single one. The full list of genes can be found in Supplementary Table 2.

**Figure 4:**
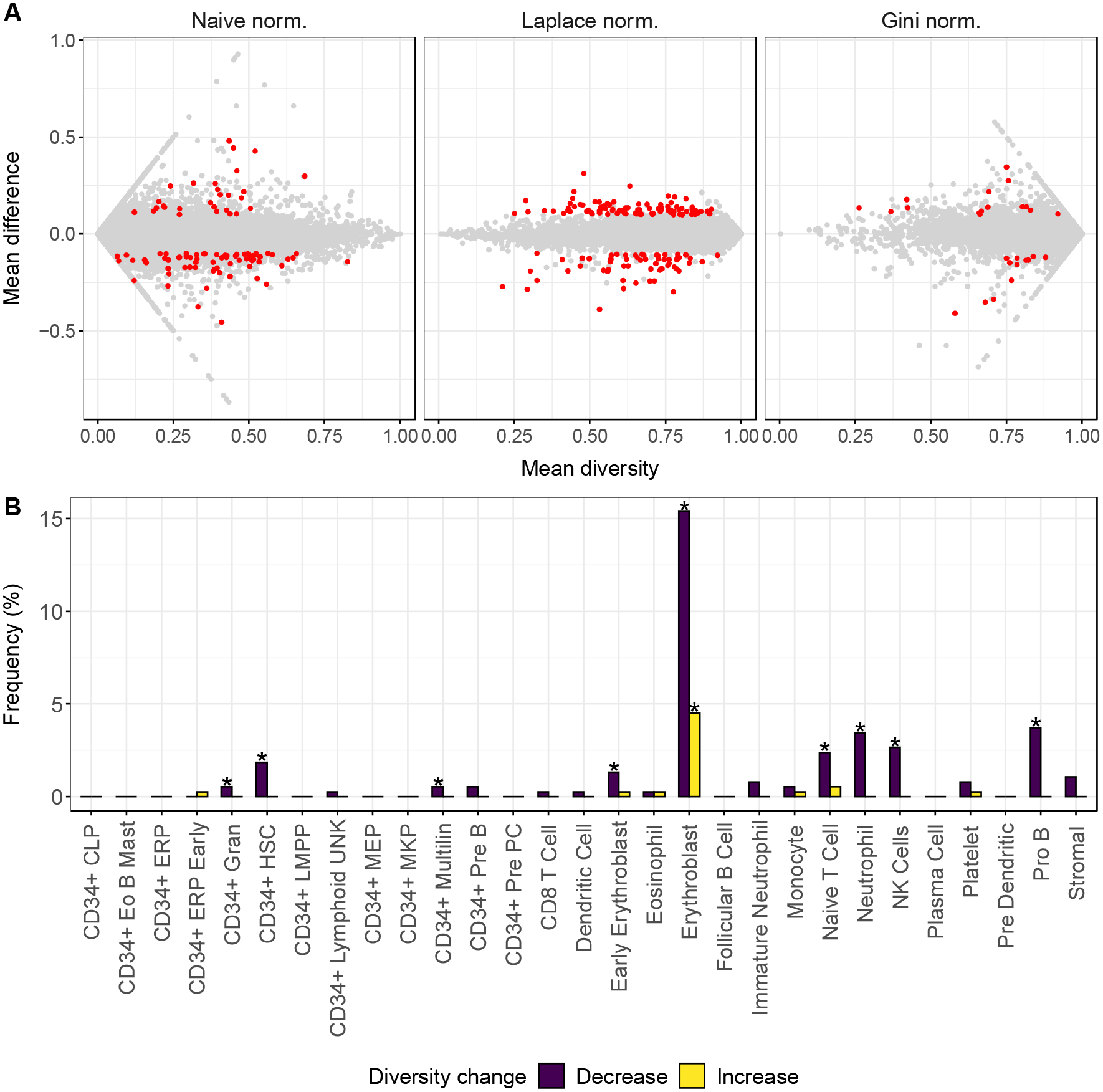
Differential diversity analysis results using MDS data. **A**) Differential diversity results comparing SF3B1 mutated and non-mutated MDS samples for 3 different diversity metrics and Salmon expression estimates. The x axis shows the mean diversity across all samples, while the y axis shows the difference of means between the SF3B1 mutated and non-mutated MDS sample groups. Each gene is a single grey dot, and significant changes are highlighted in red. Significant changes are defined as |mean difference| > 0.1 and Wilcoxon-test adjusted P-value < 0.05. **B**) Enrichment of the (Hay *et al*., 2018) marker gene sets in the differential diversity results separated by increasing or decreasing diversity between the SF3B1 mutated and non-mutated sample groups. The x axis shows the different marker gene sets, while the y axis shows the % of significantly changing genes that are falling into a specific marker gene set. Significant enrichment of a set is marked with an asterisk (Benjamini-Hochberg adjusted Fisher-test P-value < 0.05).

The 6 genes detected by all three methods as significantly changing were *ABCC5, ACOT13, DCAF16, MPC2, MRPS21* and *TMEM14C. ABCC5* is described as differentially spliced in *SF3B1* mutated breast cancer (Maguire *et al*., 2015), and *MRPS21* is described as a gene, whose upregulation is associated with poor response to azacytidine in MDS and related cancers (Belickova *et al*., 2016). We did not find a clear association to *SF3B1* mutations, cancer or hematopoiesis for the other 4 genes.

### Enrichment analysis of differential diversity genes

Finally, we chose the full list of differential diversity genes detected by the normalized naive-entropy metric, to investigate them further. We separated the 97 genes into two lists, with increased or decreased diversity in the *SF3B1* mutated samples compared to wild type, and checked their overlap with the (Hay *et al*., 2018) marker gene sets. Results are plotted on **Figure 4B**. We found several marker gene sets that were enriched in our results, with the largest enrichment in the erythroblast set. These results are consistent with the literature, as *SF3B1* mutations are known to be associated with impaired erythropoiesis (Obeng *et al*., 2016; Conte *et al*., 2015). Moreover, genes with a changing diversity of splicing isoforms might be the result of changing splicing regulation upon *SF3B1* mutations. Mutations in the yeast ortholog of *SF3B1* is known to alter the fidelity of branch site selection (Carrocci *et al*., 2017). Therefore, mutated *SF3B1* in human cells might change the splicing regulation of specific genes, leading to changes in the otherwise coordinated expression of specific splicing isoforms.

## Discussion

In this study, we have developed a package called SplicingFactory that enables the analysis of splicing isoform diversity in biological samples and between different conditions. Our tool has several benefits. It integrates into the Bioconductor R package ecosystem, and uses standard data input and output structures, facilitating analysis and integration into bioinformatics pipelines. SplicingFactory is able to process expression quantification results from a full transcriptome annotation and a large number of samples using limited hardware resources and completes in a reasonable amount of time. The tool can process any kind of expression value, although values not normalized for transcript length are not recommended. It implements 7 different diversity metrics, and methods to test differences between sample groups. As our tool provides gene and sample diversity values, it can be the basis of independent downstream analyses, not anticipated by us.

We demonstrate the usefulness of our tool by analyzing a set of CD34+ control and MDS RNA-seq samples. Most MDS patients have at least one oncogenic mutation, with splicing factor mutations occurring in more than 50% of patients (Papaemmanuil *et al*., 2015), including SF3B1, SRSF2, U2AF1 and ZRSR2. These splicing factor mutations might lead to the aberrant post-transcriptional regulation of downstream genes and the generation of oncogenic transcript isoform variants. As splicing factors are mutated early in MDS and they can be already found in normal peripheral blood, or in clonal hematopoiesis of indeterminate potential (CHIP) (Xie *et al*., 2014), they are probably responsible for increased cellular growth and clonal expansion from an early stage. However, splicing changes are generally minor and variable (Ilagan *et al*., 2015; Kim *et al*., 2015; Shirai *et al*., 2015), and do not frequently affect known MDS oncogenes or tumor suppressors (Shiozawa *et al*., 2018). Even using large-scale transcriptome datasets, complemented by targeted sequencing of key mutated genes, it seems to be difficult to point to a specific set of genes, whose transcript isoform changes are responsible for MDS development.

Mutations in spliceosome genes might act as amplifiers of splicing noise, and lead to large fluctuations in complex gene regulatory networks, altering cell fate, pathogenicity or disease penetrance (Chalancon *et al*., 2012). Initial results exist, where the authors investigated gene expression variance, showing that higher expression variability is connected to the more aggressive subtype of chronic lymphocytic leukemia (Ecker *et al*., 2015). However, the authors did not investigate splicing variance. In this study we present a list of genes, whose splicing variance is changing upon *SF3B1* mutations. Some of these genes are already known to be involved in the pathogenesis of MDS, and they are enriched for specific marker gene sets, involved in hematopoietic differentiation.

Additional analyses are needed to understand the role splicing diversity in cellular differentiation, development and disease, and we hope our tool will facilitate them.

## Availability of source code and data

The processed datasets are available in SRA with the ids SRP133442 and SRP149374. The SplicingFactory package in binary and source code format is freely available under the GPL-3.0 license from Bioconductor for the Windows, MacOS and Linux operating systems https://bioconductor.org/packages/release/bioc/html/SplicingFactory.html.

## References

Afsari, B. et al. (2018) Splice Expression Variation Analysis (SEVA) for inter-tumor heterogeneity of gene isoform usage in cancer. Bioinformatics, 34, 1859–1867.

Baralle, F.E. and Giudice, J. (2017) Alternative splicing as a regulator of development and tissue identity. Nat. Rev. Mol. Cell Biol., 18, 437–451.

Belickova, M.M. et al. (2016) Up-regulation of ribosomal genes is associated with a poor response to azacitidine in myelodysplasia and related neoplasms. Int. J. Hematol., 104, 566–573.

Van den Berge, K. et al. (2019) RNA Sequencing Data: Hitchhiker’s Guide to Expression Analysis. Annu. Rev. Biomed. Data Sci., 2, 139–173.

Carrocci, T.J. et al. (2017) SF3b1 mutations associated with myelodysplastic syndromes alter the fidelity of branchsite selection in yeast. Nucleic Acids Res., 45, 4837–4852.

Chalancon, G. et al. (2012) Interplay between gene expression noise and regulatory network architecture. Trends Genet., 28, 221–232.

Conte, S. et al. (2015) Aberrant splicing of genes involved in haemoglobin synthesis and impaired terminal erythroid maturation in SF3B1 mutated refractory anaemia with ring sideroblasts. Br. J. Haematol., 171, 478–490.

Dong, Y. et al. (2012) Anopheles NF-κB-regulated splicing factors direct pathogen-specific repertoires of the hypervariable pattern recognition receptor AgDscam. Cell Host Microbe, 12, 521–530.

Dvinge, H. et al. (2016) RNA splicing factors as oncoproteins and tumour suppressors. Nat. Rev. Cancer, 16, 413–430.

Ecker, S. et al. (2015) Higher gene expression variability in the more aggressive subtype of chronic lymphocytic leukemia. Genome Med., 7, 8.

Eling, N. et al. (2019) Challenges in measuring and understanding biological noise. Nat. Rev. Genet.

Hattori, D. et al. (2009) Robust discrimination between self and non-self neurites requires thousands of Dscam1 isoforms. Nature, 461, 644–648.

Hay, S.B. et al. (2018) The Human Cell Atlas bone marrow single-cell interactive web portal. Exp. Hematol., 68, 51–61.

Ilagan, J.O. et al. (2015) U2AF1 mutations alter splice site recognition in hematological malignancies. Genome Res., 25, 14–26.

Im, H. et al. (2018) Distinct transcriptomic and exomic abnormalities within myelodysplastic syndrome marrow cells. Leuk. Lymphoma, 59, 2952–2962.

Kim, E. et al. (2015) SRSF2 Mutations Contribute to Myelodysplasia by Mutant-Specific Effects on Exon Recognition. Cancer Cell, 27, 617–630.

Kim, M. et al. (2019) SpliceHetero: An information theoretic approach for measuring spliceomic intratumor heterogeneity from bulk tumor RNA-seq. PLoS One, 14, e0223520.

Lee, Y. and Rio, D.C. (2014) Mechanisms and Regulation of Alternative Pre-mRNA Splicing. Annu. Rev. Biochem., 84, 150317182619002.

Liberzon, A. et al. (2015) The Molecular Signatures Database Hallmark Gene Set Collection. Cell Syst., 1, 417–425.

Maguire, S.L. et al. (2015) SF3B1 mutations constitute a novel therapeutic target in breast cancer. J. Pathol., 235, 571–580.

Monlong, J. et al. (2014) Identification of genetic variants associated with alternative splicing using sQTLseekeR. Nat. Commun., 5, 4698.

Obeng, E.A. et al. (2016) Physiologic Expression of Sf3b1K700E Causes Impaired Erythropoiesis, Aberrant Splicing, and Sensitivity to Therapeutic Spliceosome Modulation. Cancer Cell, 30, 404–417.

Papaemmanuil, E. et al. (2015) Clinical and biological implications of driver mutations in myelodysplastic syndromes. Blood, 122, 3616–3628.

Pellagatti, A. et al. (2018) Impact of spliceosome mutations on RNA splicing in myelodysplasia: dysregulated genes/pathways and clinical associations. Blood, 132, 1225–1240.

Ritchie, W. et al. (2008) Entropy measures quantify global splicing disorders in cancer. PLOS Comput. Biol., 4, e1000011.

Robinson, M.D. et al. (2010) edgeR: a Bioconductor package for differential expression analysis of digital gene expression data. Bioinformatics, 26, 139–40.

Schuh, L. et al. (2020) Gene Networks with Transcriptional Bursting Recapitulate Rare Transient Coordinated High Expression States in Cancer. Cell Syst., 10, 363-378.e12.

Scotti, M.M. and Swanson, M.S. (2015) RNA mis-splicing in disease. Nat. Rev. Genet., 17, 19–32.

Seiler, M. et al. (2018) Somatic Mutational Landscape of Splicing Factor Genes and Their Functional Consequences across 33 Cancer Types. Cell Rep., 23, 282–296.

Shiozawa, Y. et al. (2018) Aberrant splicing and defective mRNA production induced by somatic spliceosome mutations in myelodysplasia. Nat. Commun., 9, 3649.

Shirai, C.L. et al. (2015) Mutant U2AF1 Expression Alters Hematopoiesis and Pre-mRNA Splicing In Vivo. Cancer Cell, 27, 631–643.

Soneson, C. et al. (2016) Differential analyses for RNA-seq: Transcript-level estimates improve gene-level inferences. F1000Research, 4, 1521.

Sterne-Weiler, T. et al. (2018) Efficient and Accurate Quantitative Profiling of Alternative Splicing Patterns of Any Complexity on a Laptop. Mol. Cell, 72, 187–200.

Wan, Y. and Larson, D.R. (2018) Splicing heterogeneity: separating signal from noise. Genome Biol., 19, 86.

Xie, M. et al. (2014) Age-related mutations associated with clonal hematopoietic expansion and malignancies. Nat. Med., 20, 1472–1478.

Zambelli, F. et al. (2018) RNentropy: An entropy-based tool for the detection of significant variation of gene expression across multiple RNA-Seq experiments. Nucleic Acids Res., 46, e46.

